# Prostaglandin E2 is a Negative Regulator of Fibroadipogenic Progenitor Differentiation in Traumatically Denervated Skeletal Muscle

**DOI:** 10.64898/2026.01.17.699776

**Authors:** Christina Doherty, Monika Lodyga, Judy Correa, Caterina Di Ciano-Oliveria, Pamela J Plant, James Bain, Jane Batt

## Abstract

**Background:** Peripheral nerve trauma denervates skeletal muscle resulting in paralysis and atrophy that is reversible if timely reinnervation occurs, due to its regenerative capacity. If reinnervation is delayed muscle’s regenerative ability is exhausted and resident fibroadipogenic progenitors (FAPs) differentiate into adipocytes and fibroblasts that replace muscle with non-contractile fibrotic tissue and fat, resulting in physical disability. Prostaglandin E2 (PGE2) inhibits adipogenesis and fibrosis in other tissues. We determined whether PGE2 could inhibit fibro-fatty degradation of long-term denervated muscle.

**Methods:** We utilized the rat tibial nerve transection model, denervating the gastrocnemius and selected a 5 week post-denervation time point to represent short-term muscle denervation injury (reversible with reinnervation), and 12 weeks to represent sustained, irreversible injury. Gastrocnemius FAPs were isolated via FACS and grown in culture to assess endogenous PGE2 production and the proliferative and differentiation response to exogenous PGE2. We evaluated transcript and protein expression of PGE2 synthesizing enzyme PTGS2, PGE2 degrading enzyme 15-PGDH and markers of proliferation, adipogenesis and fibrogenesis using RT-qPCR, immunofluorescence and SDS-PAGE/Western blotting. Paracrine impact of FAPs produced PGE2 was assessed by treating C2C12 myoblasts with FAPs conditioned media.

**Results:** Transcript expression of PTGS2 was increased and 15-PGDH decreased (4.37±2.63 and -3.06±0.85 fold change respectively, p<0.05) in 5 week, but not 12 week denervated gastrocnemius, consistent with increased PGE2 production in 5 week denervated muscle. Similarly, PTGS2 transcript levels were significantly increased (2.58±0.33 fold change, p<0.05) and 15-PGDH decreased (-5.24±3.19 fold change, p<0.05) in FAPs isolated from 5 week, but not 12 week denervated muscle, demonstrating that FAPs are a source of PGE2 in short-term denervated muscle. 16,16-dimethyl PGE2 did not impact naive FAPs in vitro proliferation, but significantly inhibited their differentiation as demonstrated by 88.9%, 82.3% and 94.2% decreases in FAPs expression of adipogenic marker perilipin-1, fibrogenic marker α-smooth muscle actin (α-SMA) and lipid content respectively, mediated via PGE2 binding to the FAPs EP4 receptor. FAPs isolated from 12 week denervated muscle demonstrated increased adipogenesis and fibrogenesis vs. naive FAPs (perilipin-1 and α-SMA 7.93±2.96 and 2.00±0.33 fold increase respectively, p<0.05) and remained fully susceptible to PGE2 inhibition of fibro-adipogenic differentiation. Conditioned media from FAPs derived from 5 week, but not 12 week, denervated gastrocnemius stimulated C2C12 myoblast proliferation which was prevented by EP4 blockade.

**Conclusions:** PGE2 is identified as a novel negative regulator of FAPs differentiation in traumatically denervated muscle, suggesting the therapeutic potential of PGE2 to prevent fibro-fatty degradation of long-term denervated muscle awaiting reinnervation.

## INTRODUCTION

Peripheral nerve injuries acutely compromise physical function due to denervation-induced skeletal muscle paralysis and rapid atrophy. While the peripheral nerve downstream of a transection injury dies, the nerve will regrow from its upstream stump to re-innervate its target muscle(s). Skeletal muscle has excellent regenerative potential and will remain viable and fully receptive to reinnervation with restoration of mass and function provided that the regenerating nerve reaches the muscle in a timely fashion [1]. In contrast, with sustained denervation the regenerative capacity of muscle fails, and it is replaced by non contractile fibrotic tissue and fat [1–6]. Peripheral nerves regenerate slowly at a rate of 1 to 3 mm/day, so that proximal limb injuries, (e.g. upper arm, brachial plexus) leave the distal target muscles (e.g. hand) denervated for a year or longer [7], by which time muscle is replaced by fibro-fatty tissues resulting in permanent physical functional disability.

Traumatic muscle denervation injuries occur predominantly in motor vehicle and workplace accidents impairing quality of life and imparting significant societal costs due to increased health resource utilization, and lost productivity [8–11]. Neonatal brachial plexus palsy, which results from severing of nerves in the brachial plexus during traumatic vaginal birth due to shoulder dystocia, occurs in 1.24 per 1000 live births in the Western world [12]. Thirty percent of these otherwise healthy infants will have lifelong arm and hand functional deficits. There are currently no pharmacologic therapies to sustain long term denervated muscle awaiting reinnervation. Surgical intervention, with the aim of enabling rapid muscle reinnervation, is the standard of care [13]. The nerve is surgically repaired at the injury site to allow the downstream portion to serve as a conduit during regeneration, thereby enhancing quality muscle reinnervation and restoration of function. However, with long nerve transit distances, simple nerve repair becomes inadequate due to excessive time to reinnervation. Other operative approaches are undertaken, such as nerve transfer, which compromises a less vital muscle in favour of the denervated muscle, to achieve the best overall functional outcome for the limb.

FAPs are resident muscle progenitor cells with the potential to differentiae into adipocytes and fibroblasts [14,15]. With acute muscle injury, FAPs proliferate to produce an environment favourable for muscle satellite (stem) cell proliferation and muscle regeneration [16,17]. In contrast, with sustained denervation or chronic muscle pathologies, FAPs undergo a switch from a muscle pro-regenerative role to a phenotype with a markedly increased propensity to fibro-adipogenic differentiation, replacing muscle with fat and fibrotic connective tissue [15,18,19]. The mechanisms by which disruption of the neuromuscular junction (NMJ) and electrical silencing of muscle following denervation impact FAPs, as well as the external factors directly controlled by the intact nerve that regulate the FAPs phenotypic switch, remain unclear.

PGE2 is a bioactive lipid that is well known to inhibit fibrosis in many tissues (e.g. kidney, heart) via signalling through its EP2 and predominantly EP4 receptors [20–22]. Inhibition of PGE2 degradation in an in vivo model of direct skeletal muscle injury, mitigated fibrosis [23]. PGE2 muscle expression is significantly increased in rodent models of short-term denervation (e.g. 1 to 2 weeks) [24,25] when post-denervation muscle sequelae remain fully reversible with re-innervation. PGE2 additionally promotes vasculogenesis and satellite cell proliferation and differentiation to positively regulate muscle regenerative/hyperplastic growth [26,27]. Despite its critical role of regulating fibrosis in other organs and positive contributions to the regeneration of skeletal muscle, its impact on FAPs is unknown and its role in the regulation of FAPs mediated fibro-fatty degradation of long-term denervated skeletal muscle has not been studied.

We speculated that PGE2 may be a key regulator of the phenotypic switch of FAPs from their pro-regenerative to pro-fibrotic/adipogenic form in muscle with sustained denervation. In this study we utilized the rat tibial nerve transection model, with its validated time epochs of denervation-induced reversible vs irreversible muscle injury with fibrosis and fat infiltration [1,2], to determine the role of PGE2 in FAPs biology, and its therapeutical potential to sustain denervated muscle awaiting reinnervation.

## MATERIALS AND METHODS

### Animal Model – Tibial Nerve Transection and Gastrocnemius Denervation

The study was conducted in accordance with the guidelines of St. Michael’s Hospital, Unity Health Toronto Animal Care Committee (ACC protocol 220). Female adult Lewis rats weighing 225 g (Charles River Laboratories, Wilmington, MA, USA) were subjected to tibial nerve transection to denervate the gastrocnemius. Briefly, the rats were maintained at a surgical plane of anesthesia with 2.5% isoflurane inhalation. One hindlimb was shaved and disinfected with poviodine followed by 70% alcohol. A 1 cm incision was made on the lateral thigh from the sciatic notch to the knee, and the biceps femoris muscle was bisected with blunt-end scissors, revealing the sciatic nerve and its tibial, peroneal, and sural branches. The tibial nerve was transected 10 mm from its gastrocnemius insertion site. In the animals allocated to a denervation cohort, the proximal end of the transected tibial nerve was sutured back to the anterior surface of the biceps femoris muscle to prevent spontaneous gastrocnemius reinnervation, using an 8-0 nylon suture. In the animals allocated to the immediate repair cohort, the transected tibial nerve was immediately re-anastomosed with a 10-0 nylon suture. Hemostasis was ensured prior to closing the skin defect with 4-0 vicryl. The contralateral hindlimb remained un-operated, providing a healthy gastrocnemius muscle as an internal control for each animal.

The rats remained free roaming, with standard bedding and chow post-operatively. At 5 or 12 weeks following tibial nerve transection the rats were anesthetized and sacrificed by a T-61 intracardiac injection. The gastrocnemius muscle was immediately atraumatically dissected from the experimental and contralateral (control) hindlimbs, and each muscle was weighed and divided for downstream assays. Gastrocnemius muscle was flash frozen in liquid nitrogen for RNA and protein extraction, frozen in liquid nitrogen cooled isopentane for immunofluorescent analysis, or placed in ice-cold saline for immediate processing via flow cytometry/FACS. At minimum, 3 independent rats were used per assay. The tibial nerve transection model is extensively utilized and well-validated (example references listed in Supplemental Materials).

### Flow Cytometry/FACS

FAPs were identified and isolated via flow cytometry/fluorescence activated cell sorting (FACS) as previously described [18]. Briefly, a single cell suspension was generated from the gastrocnemius by mechanical and enzymatic digestion. The isolated cells were immuno-labeled with fluorophore-conjugated primary antibodies CD31::FITC (Abcam ab33858, Waltham, MA, USA, an endothelial cell marker), CD45::FITC (Biolegend 202205, San Diego, CA, USA, a hematopoietic cell marker), VCAM-1::PE (Biolegend 200403 San Diego, CA, USA, a satellite cell marker), and Sca-1 (Millipore Sigma ab4336, Oakville, ON, Canada, a FAPs marker) detected with goat anti-rabbit Alexa Fluor 647 (Invitrogen, A21244, Burlington, ON, Canada). SYTOX blue staining (Invitrogen, S34857, Burlington, ON, Canada) indicated cell viability. Unstained, single-stained and fluorescent-minus-one controls were used for each experiment. FAPs (CD45-/CD31-/VCAM-1-/Sca1+) were isolated from the samples for cell culture using the Aria III (BD Biosciences, Mississauga, ON, Canada).

### Cell Culture

#### FAPs

FACS isolated FAPs were plated at a density of 5000 cells/cm2 on gelatin coated (Millipore Sigma, G1890, Oakville, ON, Canada) 6, 12, or 24 well plates for protein extraction (Sarstedt, 83.3920, Numbrecht, Germany), RNA extraction (Thermo Fisher, 130185, Burlington, ON, Canada) and immunofluorescence/ORO staining (Corning, 353047, Oakville, ON, Canada) respectively, and maintained at 37C with 5% CO2 in basal growth media (79% DMEM, 20% FBS, 1% Penicillin-Streptomycin, 10% Heat Inactivated Horse Serum, 2ng/mL basic Fibroblast Growth Factor) for 7 days until confluent. To assess fibrogenesis, FAPs were left in growth media for an additional 7 days. To assess adipogenesis, FAPs were transferred at confluency to adipogenic differentiation media (79% DMEM, 20% FBS, 1% P/S, 1.25uM Dexamethasone, 0.5mM IBMX, 5uM Troglitazone, 1ug/mL Humulin R) for an additional 7 days. In both cases media changes were made every 3 days.

To passage, FAPs were grown to 70% confluency, washed with sterile Phosphate Buffered Saline (PBS) (Gibco, 10010-023, Paisley, UK), detached with trypsin (5 min, 37°C), and plated at a density of 3500 cells/cm2 in basal growth media, with media changes every 3 days. Cells were taken out to 5 passages. P0 and P1 FAPs were used for experimentation.

To evaluate the effects of PGE2 on FAPs proliferation and differentiation, 16, 16 dimethyl-PGE2 (TOCRIS, 39746-25-3, Minneapolis, MN, USA) was reconstituted in dimethyl sulfoxide (DMSO), according to the manufacturer’s specifications. PGE2 (1nM, 100nM) and/or the EP4 specific antagonist E7046 (100nM, Cayman Chemicals, Ann Arbor, MI, USA, reconstituted in DMSO as per the manufacturer specifications), were added to FAPs cultures at the time of cell seeding and at each media change.

#### Murine C2C12 myoblasts

C2C12 murine myoblasts were plated in 12 well plates in FAPs basal growth media at a density of 1000 cells/cm2 and maintained at 37°C with 5% CO2. Conditioned media collected daily from cultured FAPs, derived from denervated or control gastrocnemius muscle over 7 days, was applied neat to C2C12 myoblasts daily for 7 days, with or without DMSO vehicle negative control and EP4 receptor antagonist E7046 (100nM). PGE2 (50 or 100nM) supplementation to basal growth media served as a positive control.

### RNA extraction and RT-qPCR

mRNA was extracted from whole gastrocnemius muscle and/or FACS isolated FAPs to quantify transcript expression of markers of fibrosis, adipogenesis, PGE2 biosynthesizing and degrading enzymes, PGE2 receptors (EP1, EP2, EP3, EP4) and housekeeping genes. RT-qPCR data was analyzed using the relative comparative ΔΔCT method. Detailed experimental methods are described in Supplemental Materials. RT-qPCR primers are show in Supplementary Table 1.

### Protein Extraction, SDS/PAGE and Western Blotting

Briefly, whole gastrocnemius protein lysates and FAPs nuclear protein lysates were separated by SDS/PAGE and Western blotting performed for COX-2, EP4 and PPARγ. Chemiluminescence was detected using ClarityWestern ECL Substrate (Bio-Rad, 1705060S, Mississauga, ON, Canada) with signal acquired on a Gel Doc EZ Imager (Bio-Rad Laboratories, Mississauga, ON, Canada). Chemiluminescent signal was quantified on Image Lab 6.0.1 (Bio-Rad, Mississauga, ON, Canada). Detailed experimental methods are described in Supplemental Materials.

### Immunofluorescence and ORO staining

Cultured FAPs were immunostained for Ki67, perilipin-1 (PLIN-1), α-smooth muscle actin (α-SMA), and PPARγ, detected with fluorescent secondary antibodies, or stained with Oil Red O (ORO), to enable quantification of proliferation, adipogenic and fibrogenic differentiation. Detailed experimental methods are described in Supplemental Materials.

### Microscopy, Image Acquisition and Quantification

Cultured cells were imaged with the BioTek Cytation 5 Imaging System (Agilent, Mississauga, ON, Canada) with a SONY IMX264 CMOS camera and a 10×/0.3 Plan Fluorite objective or the Zeiss Wide Field, with an Axiocam712 mono 4096x3008 detector, and either a 10x0.45 or 40x0.6 objective. Acquisition settings were kept constant across experimental conditions for each individual stain, and 8 image fields were randomly acquired per stain. For each image nuclear counts were determined with Fiji software Version 1.54F (https://imagej.net/contribute/citing; script for the macro is in Supplemental Materials) and number of cells positive for PLIN-1, PPARγ or ORO was manually performed by a trained, blinded reviewer. PLIN-1, PPARγ or ORO percent positive cells were determined as the number of PLIN-1, PPARγ or ORO positive cells/total number of cells in the microscopic field. A minimum of 1500 cells were evaluated per experimental sample.

### Statistical Analysis

Statistical analysis was completed with GraphPad Prism (10.2.3). Shapiro-Wilks test was used to test for normal data distribution. Comparisons between the experimental conditions at each time point post-denervation were completed with t-tests, one way ANOVA or Kruskal Wallis with post-hoc multiple comparison’s test, as appropriate. Statistical significance was set at p < 0.05.

## RESULTS

### PGE2 temporal expression in denervated muscle coincides with time-dependent reversibility of denervation-induced muscle fibrosis and fat deposition

To determine if PGE2 expression remains elevated in muscle following sustained denervation injury in a timeline co-incident to the establishment of irreversible fibrosis and fat infiltration, we evaluated expression of the enzymes that produce and degrade PGE2. Arachidonic acid (AA) is metabolized sequentially by cyclo-oxygenase 2 (COX-2, encoded by the gene PTGS2) to produce the intermediary PGH2, followed by prostaglandin E synthase (PTGES) to produce PGE2 [21]. PGE2 is inactivated and degraded through oxidation by 15-hydroxyprostaglandin dehydrogenase (15-PGDH) [21]. The stages of denervation-induced muscle sequelae are well defined [1,2]. We used 5 week (5wk) denervated gastrocnemius to represent the initial stage of muscle denervation injury - the established time epoch from 0 to 2 months post-denervation in the rat when muscle sequelae remain fully reversible with reinnervation. A 12 week (12wk) post-denervation time point was selected to represent the second stage epoch from >2 to 7 months, when irreversible muscle fibrosis and fat deposition commence and receptivity to reinnervation declines. We found that PTGS2 transcript/COX-2 protein levels increased significantly in gastrocnemius muscle at 5 wks post denervation but returned to baseline levels or below with sustained denervation (12wk) (Fig. 1A,C). In contrast, transcript expression of 15-PGDH was significantly decreased at 5wks and increased at 12wks post-denervation (Fig.1B). These data suggest muscle PGE2 expression increases transiently following denervation when muscle sequelae remain reversible but subsequently declines in the absence of reinnervation, when irreversible muscle fibrosis and fat infiltration ensue.

**Figure 1.**
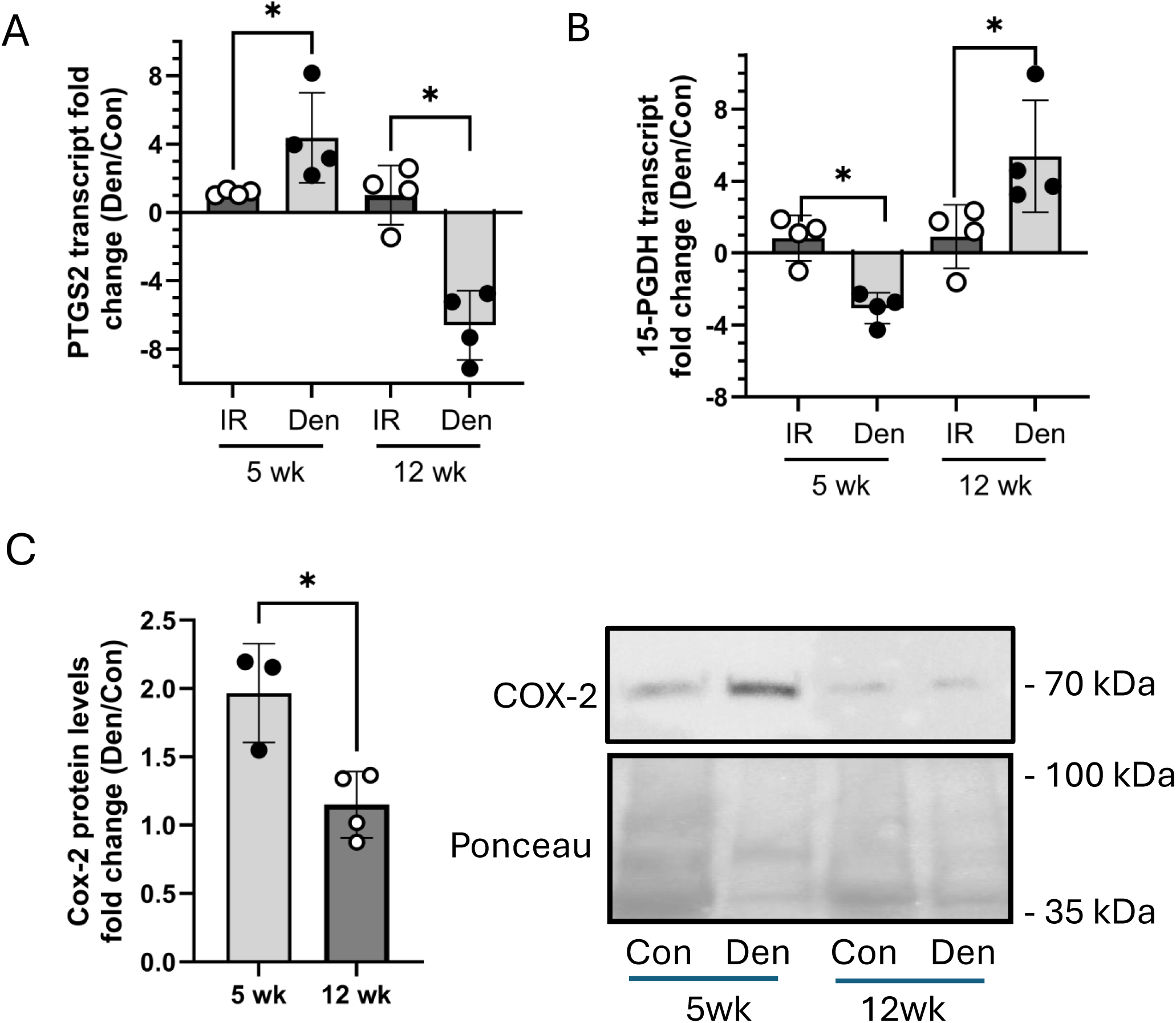
PGE2 synthesizing and degrading enzyme expression in denervated gastrocnemius. Tibial nerve transection was performed, and the gastrocnemius was either left denervated (Den) or underwent subsequent immediate repair of the nerve (IR) to reinnervate the gastrocnemius. The muscle was harvested at 5 or 12 weeks (wks) post procedure. The non-operated contralateral gastrocnemius served as internal control (Con). Gastrocnemius PTGS2 (A) and 15-PGDH (B) transcript expression was determined by RT-qPCR. HPRT and HMBS served as housekeeping genes. (C) 40ug gastrocnemius protein lysates were separated by SDS-PAGE and COX-2 protein levels determined by Western blotting, quantified by CCD camera acquisition of the chemiluminescent signal. Representative blots are shown. Ponceau staining served as loading control. Data for the experimental gastrocnemius was normalized to the control gastrocnemius for all analyses. (Data are mean +/- SD, *p<0.05)

### FAPs produce PGE2 and express PGE2 (EP) receptors

To determine if FAPs are a source of PGE2 in skeletal muscle we evaluated PTGS2, PTGES and 15-PGDH transcript expression in FAPs derived from denervated and contralateral control gastrocnemius muscle. PTGS2 mRNA was differentially expressed across the two time points, increased at 5wks and decreased at 12wks post-denervation relative to baseline expression, while 15-PGDH expression was significantly decreased at 5wks and increased at 12wks post-denervation (Fig. 2A), mirroring the changes observed in whole muscle. We also assessed the 3 isoforms of prostaglandin E synthase (PTGES1, 2 and 3) that metabolize the intermediary PGH2 to PGE2. PTGES1 and PTGES2 were significantly upregulated in FAPs isolated from short-term (5wk), but downregulated in FAPs from long term (12wk) denervated muscle. Together these data suggest that FAPs are a source of the transiently increased muscle production of PGE2 following denervation injury when muscle sequelae remain fully reversible (Stage 1), but decrease PGE2 production with sustained denervation (Stage 2).

**Figure 2.**
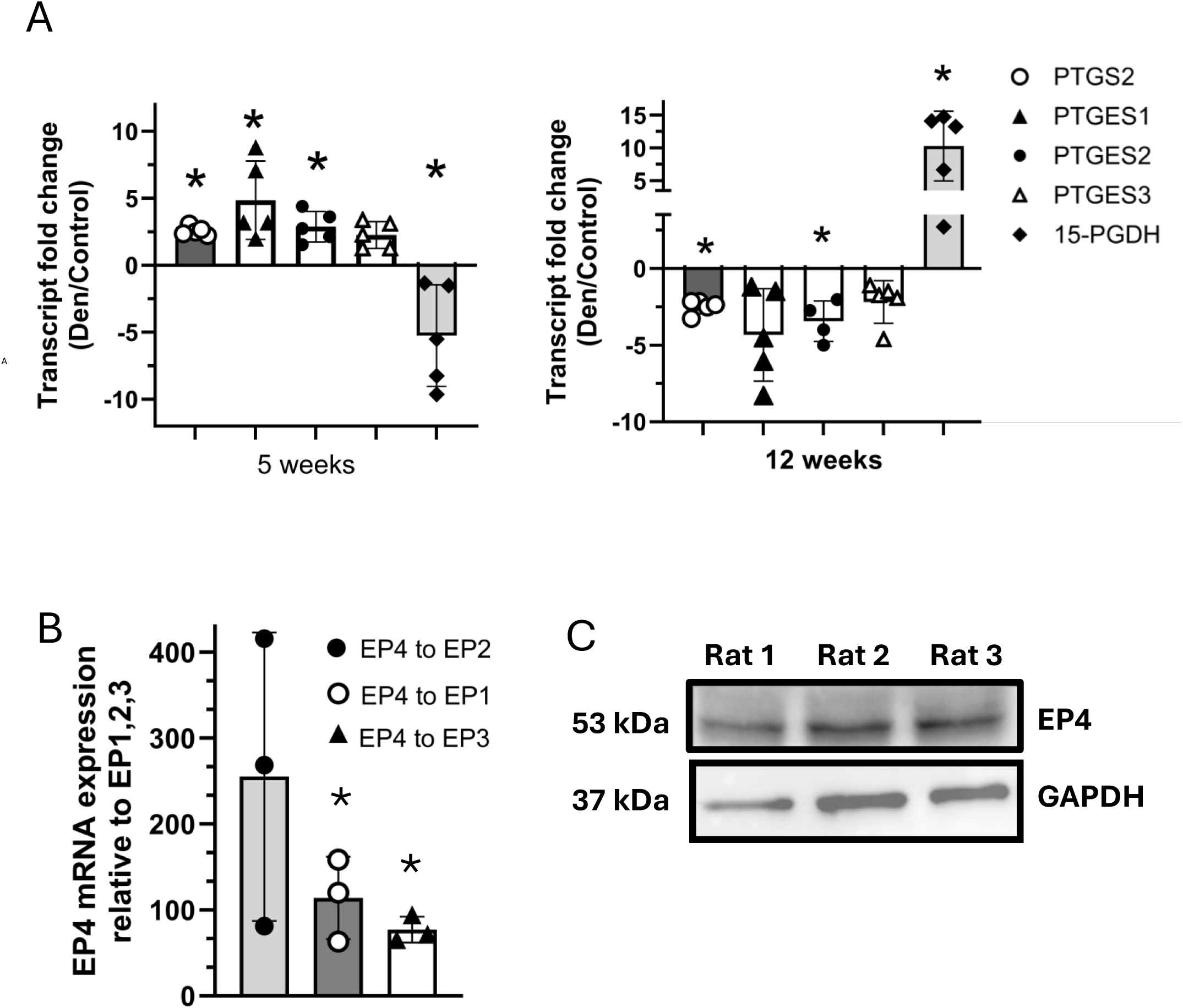
FAPs produce PGE2 and express EP receptors. (A) RT-qPCR determined fold change in expression of PGE2 synthesizing enzymes (PTGS2, PTGES1, PTGES2, PTGES3) and PGE2 degrading enzyme (15-PGDH) transcript levels in FAPs isolated from 5 week (left panel) and 12 week (right panel) denervated gastrocnemius vs contralateral control gastrocnemius, representing time epochs of reversible and irreversible denervation mediated injury respectively. FAPs increase PGE2 synthesizing enzymes and decrease 15-PGDH expression at 5 weeks, with a reciprocal pattern of expression at 12 weeks post-denervation. (B) FAPs isolated from naïve gastrocnemius express EP1, EP2, EP3 and EP4 receptor transcripts. EP4 expression is dominant. (C) 10ug FAPs protein lysate was separated by SDS-PAGE and probed for the EP4 receptor. Representative Western blots are shown. GAPDH served as loading control. (Data are mean +/-SD, *p <0.05)

PGE2 is known to bind one of four EP receptors (EP1, EP2, EP3, EP4) in its target cells to signal downstream [21,28]. To determine if FAPs might respond in an autocrine manner to the PGE2 they produce, FAPs EP receptor expression was assessed (Fig. 2 B,C). FAPs were found to express all 4 EP receptors, with EP4 being dominant at over 70-fold higher transcript expression relative to EP receptors 1 through 3.

### PGE2 inhibits FAPs adipogenic and fibrogenic differentiation

To determine if PGE2 impacts FAPs phenotype we evaluated the effect of exogenous PGE2 on FAPs proliferation and differentiation in tissue culture. Exposure of FAPs isolated from naïve gastrocnemius to exogenous PGE2 at physiologic concentrations *in vitro* did not impact FAPs proliferation (Fig. 3), but markedly mitigated both their adipogenic (Fig. 4) and fibrogenic (Fig. 5) differentiation, as determined by evaluating cellular expression of adipogenic marker PLIN-1, PPARγ (a pro-adipogenic transcription factor), cellular lipid content and fibrogenic markers α-SMA and collagen 1A1.

**Figure 3.**
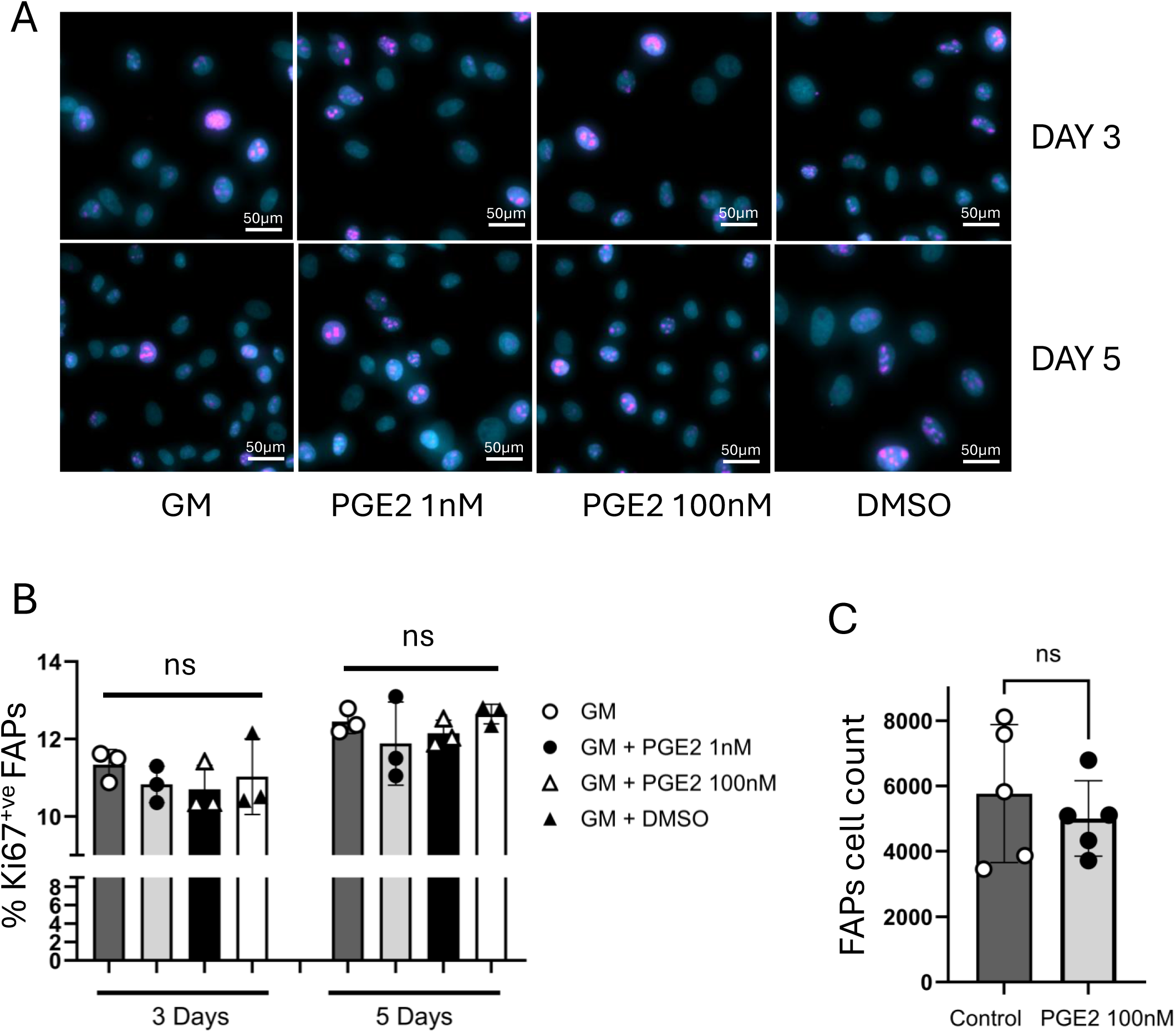
PGE2 does not impact FAPs proliferation. (A,B) PGE2 did not impact *in vitro* proliferation of FAPs derived from naïve gastrocnemius at 3 and 5 days post plating, as determined by quantification of nuclear expression of proliferation marker Ki67. Representative images of FAPs cultures immunostained for Ki67 (pink), nuclei in turquoise, are shown. (C) FAPs cell count at 7 days post plating was not impacted by PGE2 (GM = growth media, DMSO = vehicle control, Data are mean +/- S.D., ns = not significant [p≥0.05]).

**Figure 4.**
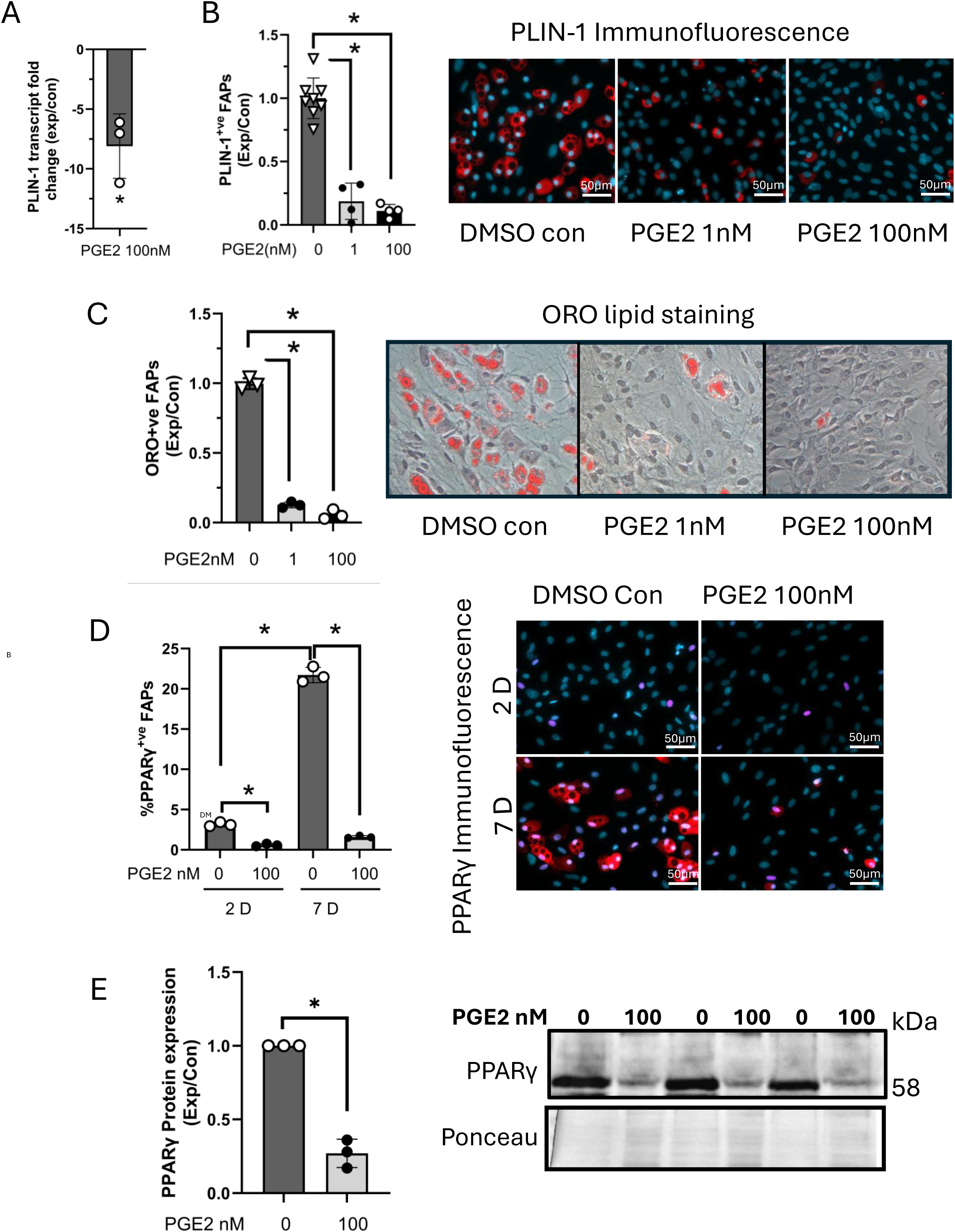
PGE2 inhibits naïve FAPs adipogenic differentiation. FAPs isolated from naive gastrocnemius muscle were cultured in adipogenic differentiation media with or without PGE2 or DMSO (vehicle negative control) and differentiation assessed by quantifying FAPs expression of the adipogenic marker perilipin-1 (PLIN-1), and lipid content at 7 days. Results in experimental FAPs cultures were normalized to control cultures (adipogenic media alone). PGE2 significantly decreased (A) PLIN-1 transcript (RT-qPCR) and (B) protein expression (immunofluorescence - PLIN-1 in red, nuclei in turquoise), and (C) lipid content (Oil Red O [ORO]; lipid stain in red). (D) PPARγ is a transcription factor required for adipogenesis. PGE2 decreased FAPs nuclear PPARγ protein expression (immunofluorescence- PPARγ in pink, PLIN-1 in red, nuclei in turquoise ) at 2 and 7 days post plating. (E) 10ug of FAPs nuclear protein lysates were separated by SDS-PAGE and immunoblotted for PPARγ, similarly showing PGE2 inhibits FAPs PPARγ expression. Ponceau staining served as loading control. Representative Western blots are shown. (Data are mean +/- S.D, *p<0/05, Exp = experimental cultures, Con = media control, D = Days).

**Figure 5.**
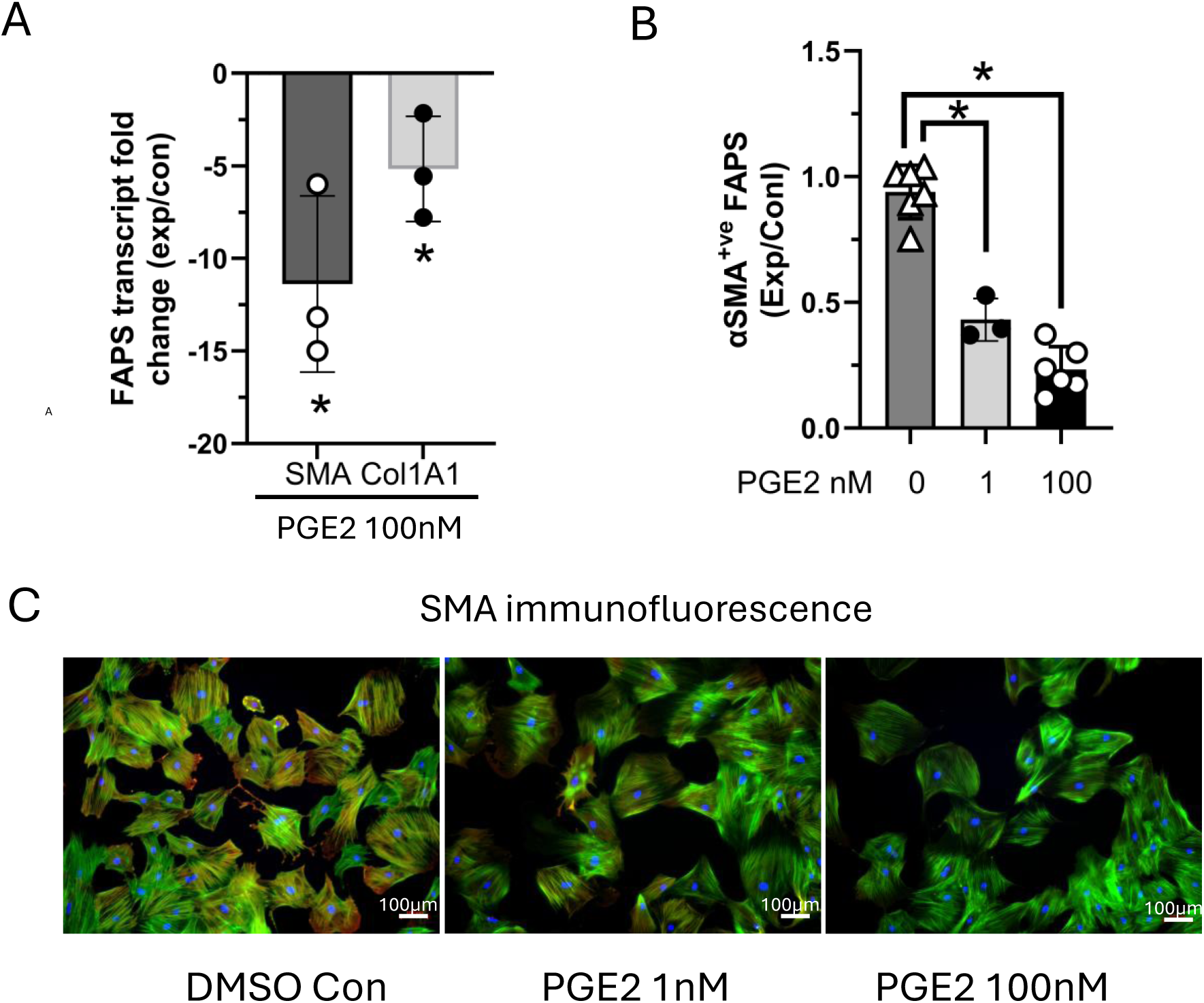
PGE2 inhibits naïve FAPs fibrogenic differentiation. FAPs isolated from naive gastrocnemius muscle were cultured in growth media with or without PGE2 or DMSO (vehicle negative control), and spontaneous fibrogenic differentiation assessed by quantifying expression of the fibrogenic marker markers Collagen 1A1 (Col-1A1) and αSMA (α-smooth muscle actin) 14 days post plating. Results in experimental FAPs cultures were normalized to control cultures (growth media alone). PGE2 decreased (A) αSMA and Col-1A1 transcript expression (RT-qPCR) and (B,C) αSMA protein expression (immunofluorescence – Phalloidin is green marking F-actin, αSMA in red, nuclei in blue). (Data are mean +/- S.D., *p<0.05, Exp = experimental culture, Con = control media.)

FAPs derived from 12 wk (Stage 2) denervated rat muscle are known to demonstrate significantly increased differentiation capacity, in keeping with the temporal switch of muscle from reversible, to progressive and irreversible fibro-fatty degradation induced by sustained denervation. We found that FAPs isolated from 12 wk denervated gastrocnemius demonstrated significantly increased adipogenic (Fig 6 A,B) and fibrogenic (Fig 6C) differentiation in culture, as previously reported. Despite this markedly increased fibro-adipogenic capacity, we also found that the 12 wk denervated FAPs remained fully susceptible to inhibition of differentiation by exogenous PGE2 (Fig. 6A,B,C).

**Figure 6.**
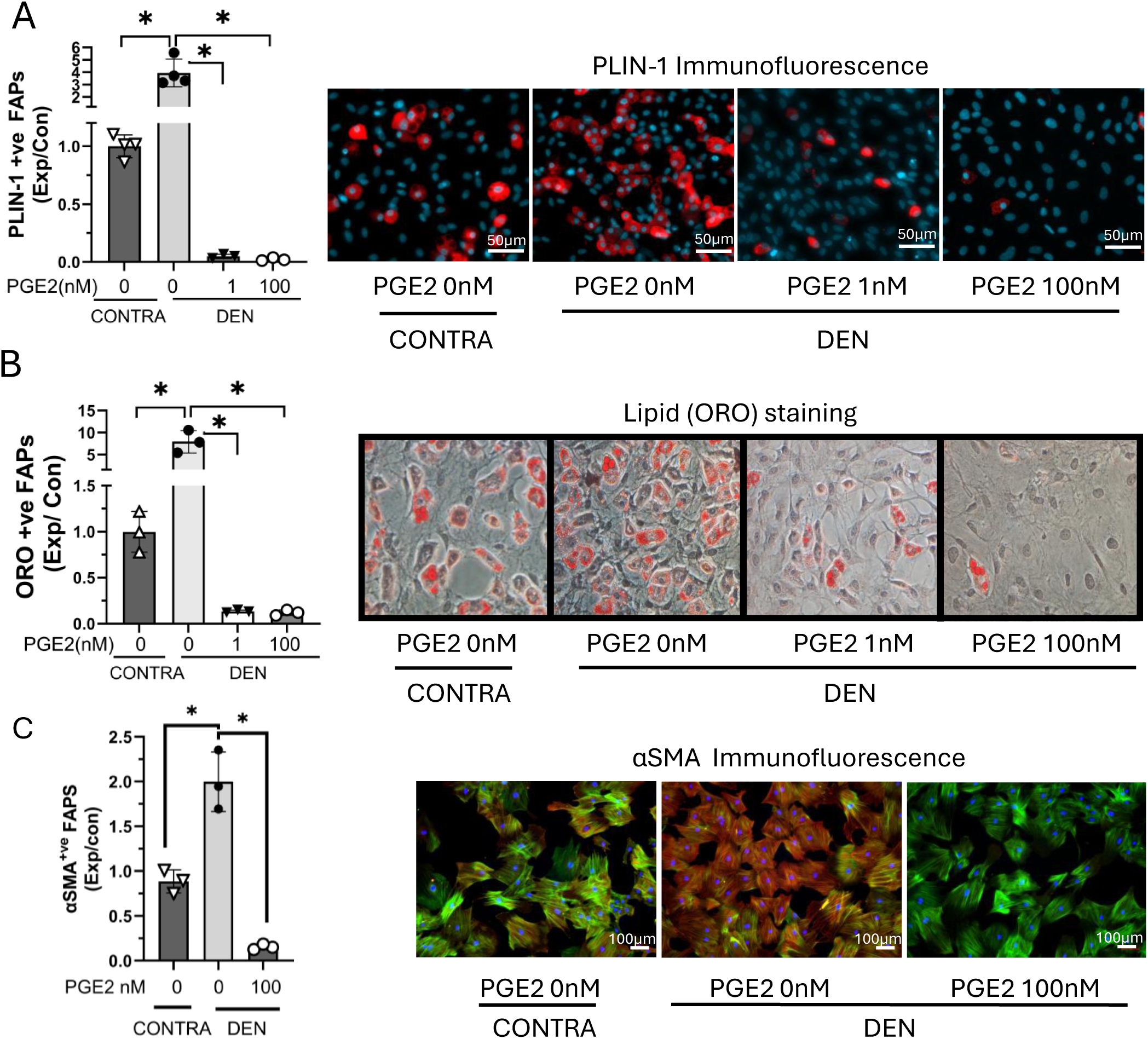
FAPs isolated from 12 week denervated gastrocnemius remain susceptible to PGE2-mediated inhibition of adipogenesis & fibrogenesis. FAPs isolated from 12 week denervated and contralateral control gastrocnemius were cultured in the appropriate media, with or without PGE2 and DMSO (vehicle negative control) and fibro-adipogenic differentiation assessed by immunostaining for relevant markers. Results in experimental FAPs cultures were normalized to control cultures (standard media). 12 week denervated FAPs demonstrate increased adipogenic differentiation as indicated by increased (A) PLIN-1 expression (immunofluorescence - PLIN-1 in red, nuclei in blue) and (B) lipid synthesis (ORO staining, lipid is red), and fibrogenic differentiation as indicated by increased (C) αSMA expression (immunofluorescence - αSMA in red, phalloidin in green, nuclei in blue) compared to FAPs isolated from the contralateral control gastrocnemius. PGE2 significantly decreased both adipogenic (A,B) and (C) fibrogenic differentiation of 12 week denervated FAPs. (DEN = denervated gastrocnemius, CONTRA = contralateral gastrocnemius, Exp = experimental culture, Con = standard media control, * p<0.05)

### PGE2 signals via the EP4 receptor to inhibit FAPs differentiation

FAPs are not a predominant muscle cell type so the total number of FAPs isolated per gastrocnemius through FACs are low. To determine whether PGE2 signals via the EP4 receptor to inhibit adipogenic and fibrogenic differentiation, naïve FAPs were passaged *in vitro* to obtain adequate cell numbers for experimentation. Retention of cellular phenotype, including differentiation capacity, EP4 receptor expression and susceptibility to PGE2 inhibition of differentiation, was confirmed for FAPs to passage 3 (Supplementary Fig. 1). Beyond P3 adipogenic and fibrogenic differentiation capacity significantly decreased. FAPs used for experimentation were P0 (freshly sorted) and/or P1.

The EP4 receptor specific antagonist, E7046 [29] did not impact FAPs basal adipogenic or fibrogenic differentiation (Fig. 7). However, the EP4 antagonist prevented PGE2 meditated inhibition of FAPs adipogenic and fibrogenic differentiation, suggesting that PGE2 signals solely via the EP4 receptor to block FAPs differentiation.

**Figure 7.**
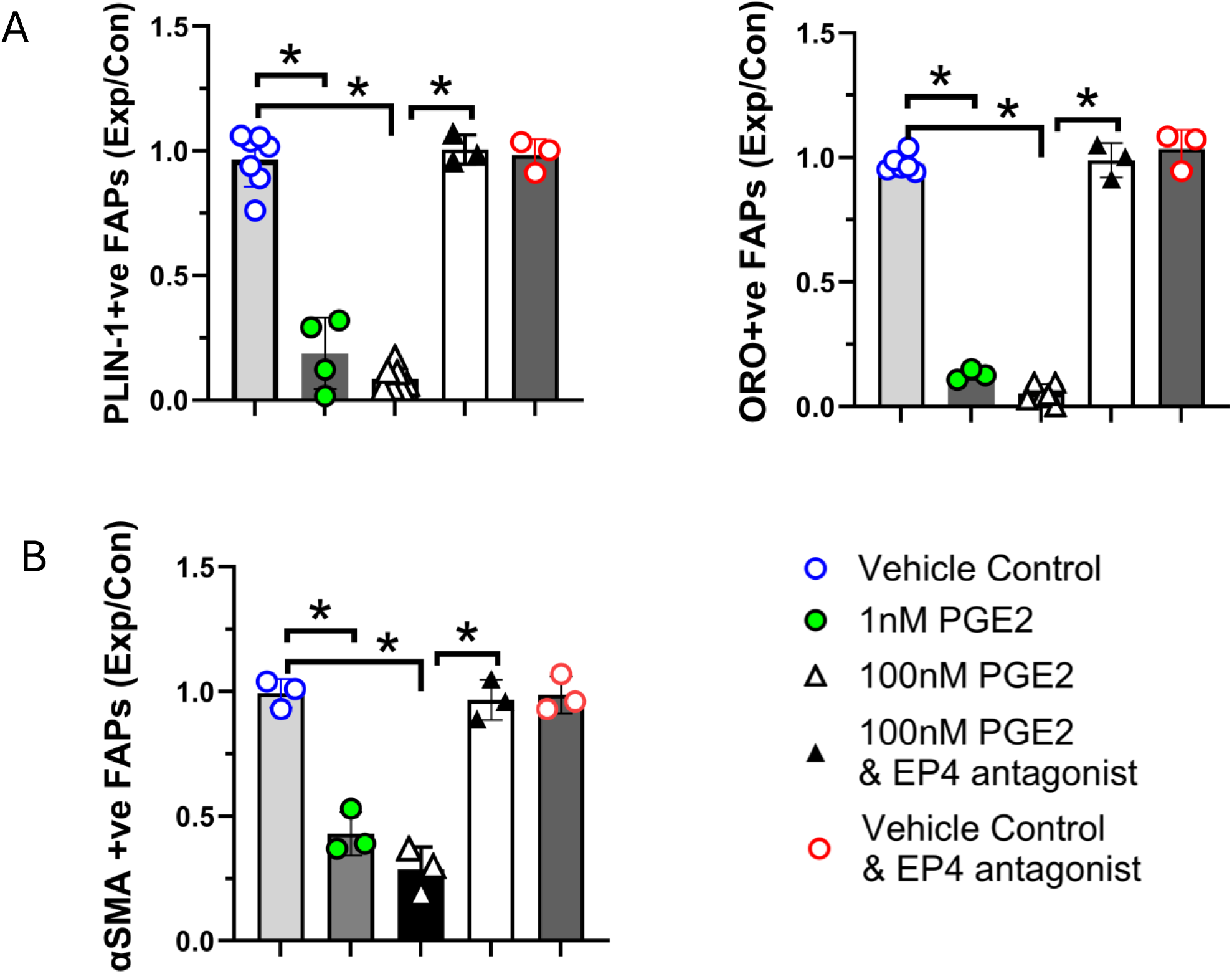
PGE2 signals via EP4 to inhibit FAPs differentiation. Naïve FAPs were differentiated in the appropriate media for 7 days with or without PGE2 and the EP4 specific antagonist E7406. DMSO served as vehicle only negative control. PGE2 inhibition of FAPs adipogenic differentiation, as determined by quantification of PLIN-1 immunostaining (left panel) and lipid synthesis (ORO staining, right panel), was prevented by the EP4 receptor antagonist E7406. Similarly, PGE2 mediated inhibition of FAPs fibrogenic differentiation was blocked by E7046, determined by quantification of αSMA immunostaining. (Data are mean+/- SD, *p<0.05, exp = experimental condition, control = control media).

### Paracrine activity of PGE2 produced by FAPs derived from short term denervated muscle stimulates C2C12 myoblast proliferation

Satellite cells and myoblasts increase proliferation substantially in the transient regenerative effort that occurs in the first stage post-muscle denervation, but this response wanes with sustained denervation and the myogenic progenitor numbers decline. [1,2,30]. Prostaglandin E2 is known to stimulate satellite cell and myoblast proliferation via the EP4 receptor [26,27]. To determine if FAPs generated PGE2 induced myoblast proliferation we treated murine C2C12 myoblasts with FAPs conditioned media in culture. We demonstrated EP4 receptor expression in C2C12 myoblasts (Supplementary Fig.2). Conditioned media from FAPs derived from 5 wk denervated gastrocnemius significantly increased C2C12 myoblast proliferation (Fig 8A) and cell count (Fig 8B), which was completely inhibited by the EP4 receptor specific antagonist, E7046. In contrast, conditioned media from 12 wk denervated FAPs had no impact on C2C12 proliferation (Fig 8A) or cell count (Fig. 8B). Together these data suggest that PGE2 produced by FAPs in short term, Stage 1 denervated muscle acts in a paracrine fashion to induce myoblast proliferation. With sustained denervation (Stage 2) however, FAPs PGE2 production and paracrine support of myoblast proliferation ceases.

**Figure 8.**
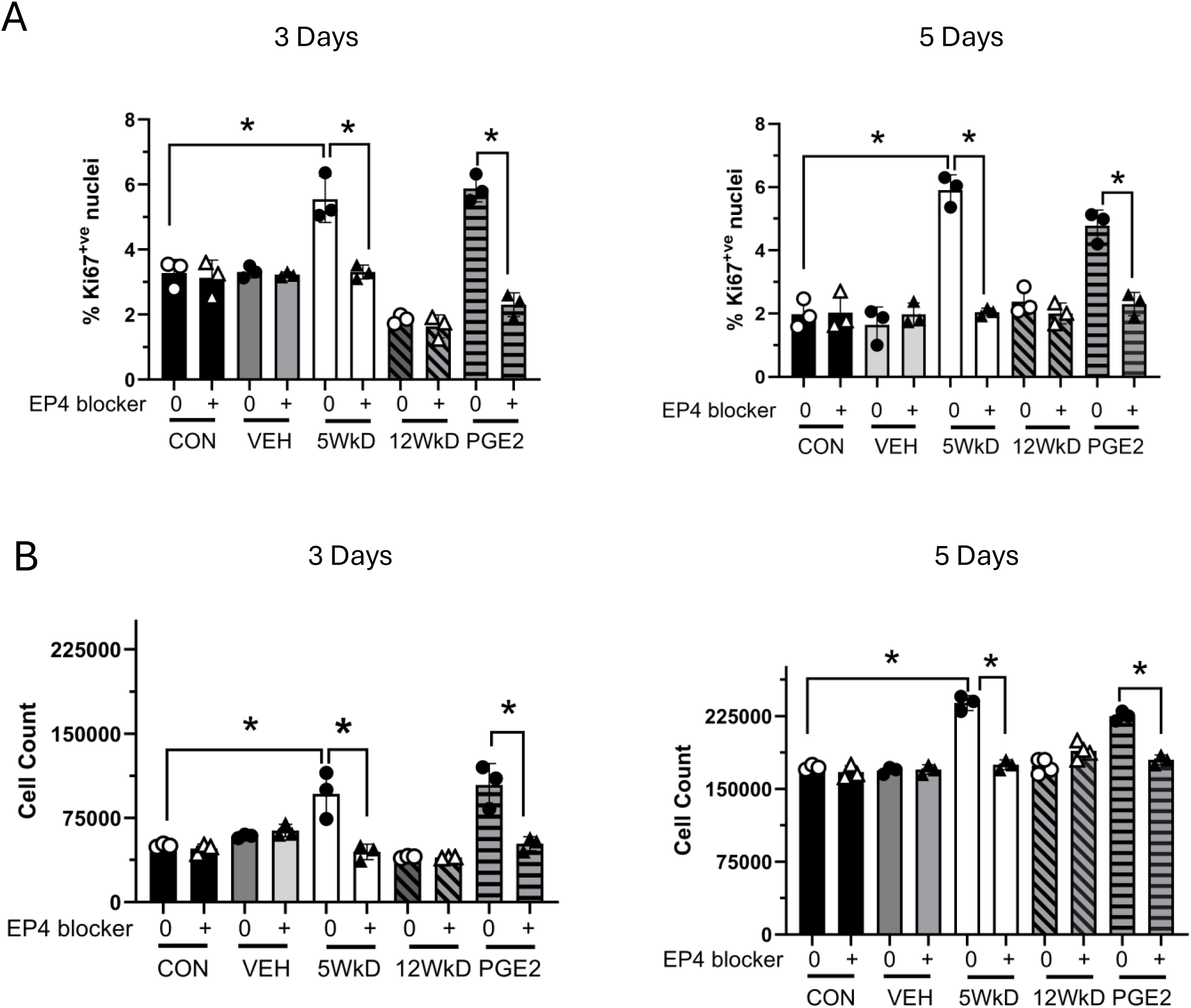
FAPs generated PGE2 stimulates C2C12 myoblast proliferation in paracrine fashion. (A) Conditioned media from 5 week denervated (5WkD), but not 12 week denervated (12WkD) FAPs, induced C2C12 proliferation as determined by (A) quantification of C2C12 nuclear expression of proliferation marker Ki67 and (B) cell count after 3 days (left panels) and 5 days (right panels) in culture. EP4 specific receptor antagonist E7046 (100nM; EP4 blocker) prevented the C2C12 proliferative response to conditioned media from 5 week denervated FAPs. DMSO (vehicle only, VEH) and conditioned media from contralateral gastrocnemius (CON) were negative controls. 16,16, dimethyPGE2 (100nM; PGE2) was positive control.

## DISCUSSION

Limb trauma with peripheral nerve injury occurs globally in working age and young populations and can result in permanent physical functional deficits because there are no means by which to sustain long-term denervated muscle awaiting reinnervation. In this study we demonstrate that PGE2 markedly inhibits FAPs fibro-adipogenic differentiation, suggesting its potential as a therapeutic option to prevent the irreversible fibrosis and fat infiltration that occurs in long-term traumatically denervated muscle. Our data support a model whereby FAPs are an early key source of PGE2 in muscle post denervation, which prevents FAPs differentiation in an autocrine feedback loop and concomitantly promotes myoblast expansion in a paracrine fashion. However, with sustained muscle denervation FAPs PGE2 production ceases, which enables FAPs persistence, fibro-adipogenic differentiation and ultimately, the irreversible muscle fibrosis and fatty degradation of long-term denervated muscle.

Although FAPs are a unique muscle resident progenitor population, they are not a homogeneous population [15,31,32]. FAPs are identified using flow cytometry and single cell RNAseq, with an array of standard positive and negative selection markers and transcriptomic signatures. Changes in expression levels of these makers, the emergence of others, and transcriptomic signature alterations in varied models of muscle injury and pathologies emphasize that the biologic role and behaviour of FAPs is highly context dependent, influenced by their baseline abundance, cellular microenvironment, muscle type, and the insult applied. Thus, the experimental model is of critical importance when delineating the cellular mechanisms underpinning FAPs regulation of pathologic processes. Denervation of the hind limb musculature of the rat has been extensively studied and has delineated well characterized time epochs that fully recapitulate the post-denervation muscle sequelae experienced in humans and the potential for reversal with reinnervation, albeit in a faster timeline [1,2]. The tibial nerve transection model specifically utilized here is extensively employed in study of the pathogenesis and treatment of denervation induced muscle injury.

Much research on FAPs biology has been conducted using models of injury that induce muscle inflammation, ranging from hyperacute insults such as myotoxin/chemical injury to chronic models of muscle fibrogenesis and adipogenesis associated with acquired obesity and diabetes, and inherent genetic anomalies (e.g. muscular dystrophies). The inflammatory infiltrate in muscle in these models can be marked, and in the case of direct myotoxin or chemical injury, can be supraphysiologic. With such acute inflammation various mediators such as interleukins 4 and 13 drive the increase in FAPs numbers, with a secretome that supports myoblast proliferation [33]. As the injurious insult resolves, FAPs undergo apoptosis and decrease to baseline numbers [34]. However, with chronic injury and inflammation FAPs are induced to persist and differentiate, resulting in muscle fibrosis and fat deposition [35].

In contrast, we and others have shown that there is no, to minimal, inflammatory response with muscle denervation acutely and resolution of inflammation with sustained denervation [14,18], suggesting that FAPs persistence and increased fibro-adipogenic differentiation is induced by other external cues. It is of interest that COX-2 activation occurs in inflammatory conditions and that PGE2 itself, produced by various cell types (e.g. neutrophils, eosinophils), can act as a pro-inflammatory mediator. We show however, that in the context of muscle denervation injury, FAPs themselves are the critical source of PGE2, stimulated to produce the biolipid early post-denervation with subsequent loss of production if denervation persists. Moreover, given that our serial assessment of COX-2 and PGDH-15 expression in whole muscle displayed a pattern in keeping with decreased PGE2 production in the 12 wk denervated gastrocnemius, it suggests that PGE2 production by other muscle resident cell types (e.g. satellite cells, endothelial cells) during sustained denervation, is insufficient to maintain FAPs quiescence and prevent their differentiation. This not surprising, considering the known satellite cell and vascular loss that occurs concomitant to progressive fibro-fatty degradation of long term-term denervated muscle [1].

Our finding that PGE2 is a potent inhibitor of FAPs adipogenic differentiation is in keeping with extensive literature demonstrating the negative impact of PGE2 on adipogenesis [28,36]. At the molecular level, adipogenesis is mainly regulated by PPARγ and CCAAT-enhancer-binding proteins (C/EBPs). PGE2 has been shown to strongly suppress the early phase of adipocyte differentiation by enhancing its own production via receptor-mediated elevation of adipocyte expression of COX-2, and additionally via suppression of nuclear PPAR*γ* expression [28,36].

While we also demonstrate profound inhibition of FAPs fibrogenic differentiation by PGE2, in contrast to adipogenesis, the reported impact of PGE2 on tissue fibrosis has been found to be variable. The inconsistent outcomes reported in the literature appear to derive from organ and model specific differences, including the dominant EP receptors and the downstream signalling networks engaged. Downstream signalling mainly mediated via the PGE2 EP2 or EP4 receptors are known to inhibit the proliferation, transformation and ECM production of fibroblasts in kidney, via inhibition of the TGFβ/SMAD signalling pathway [20,21]. EP4 activation inhibits cardiac fibrosis via PKA signalling [22]. In skeletal muscle specifically, fibrosis was mitigated in a direct injury model of the tibialis anterior muscle by using SW033291, a 15-PGDH inhibitor [23]. However, PGE2 engagement of EP1/EP3 receptors in kidney and heart has conversely been shown to promote fibrosis, signalling via the MAP kinase or ERK networks [21, 37]. Thus, evaluation of PGE2 impact is context and cell-type specific, re-iterating that model selection, to represent the pathologic process of interest, is paramount. The significant predominance of EP4 over EP1, EP2, and EP3 receptors that we observed in FAPs is in keeping with the cardiac and renal literature where PGE2 signalling, via EP4, mitigates fibrogenic differentiation.

We have also demonstrated that the favourable impact of PGE2 in denervated muscle is extended beyond FAPs. The EP4 receptor dependent proliferative response of myoblasts to the conditioned media from FAPS derived from 5 wk denervated muscle suggests that FAPs generated PGE2 supports muscle’s regenerative capacity in the early phases following nerve injury. However, as time progresses, the loss of FAPs PGE2 production depletes paracrine support to sustain muscle’s myogenic cells and its regenerative potential. Concomitantly, the loss of PGE2 mediated autocrine stimulation of FAPs permits their fibrogenic and adipogenic differentiation.

Importantly, we have also demonstrated that the susceptibility of FAPs to PGE2 mediated inhibition of differentiation is retained, regardless of the FAPS phenotype. Sca-1 is a positive selection maker of FAPs, and others have shown that Sca-1 micro-heterogeneity impacts the fate decision of dystrophic FAPs; cells with increased Sca-1 expression demonstrated markedly enhanced adipogenic differentiation in MDX mice [14,34]. We have similarly previously reported that the proportion of high Sca-1 expressing FAPs significantly increases from less than 1% in a healthy gastrocnemius FAPs population, to 50% and greater in long term denervated muscle (12 and 14 wks post-denervation), and that these long-term denervated FAPs demonstrate highly increased propensity to fibro-adipogenic differentiation [18,30]. Here we show that PGE2 continues to fully suppress both the fibrogenic and adipogenic differentiation of FAPS derived from muscle experiencing sustained denervation.

While we have identified the production of PGE2 by FAPs as being regulated by innervation, we have not identified the upstream factors responsible for this impact. There has been extensive research evaluating the intracellular signalling the drives FAPs proliferation and differentiation in various muscle injury models, but outside of infiltrating inflammatory cells external factors remain largely undefined. It has been previously reported that FAPs closely interact with Schwann cells in the neuromuscular junction and muscle-nerve associated niches, that FAPs can impact Schwann cell homeostasis and that there is crosstalk between glial cells and FAPs [14,35,38,39]. How these nerve-FAPs interactions might regulate FAPs COX-2 activity and PGE2 production, is an area for future study.

This study has limitations. We did not undertake metabolomic assays to directly demonstrate PGE2 production by FAPs, but showed instead indirect evidence of PGE2 production, by evaluating the level of both PGE2 synthesizing and degrading enzymes (protein and/or transcript expression) in both whole muscle and FAPs. Moreover, conditioned media from short term (5wk) denervated FAPs, where COX-2, PTGES-1, PTGES-2 and PGDH-15 transcript and/or protein levels indicated increased FAPs PGE2 production, was able to induce C2C12 proliferation that was completely blocked with a EP4 receptor specific antagonist, again supporting FAPs PGE2 production.

As noted previously, FAPs are highly heterogeneous, and functional variability across subpopulations appears to be influenced by multiple factors, including injury specific cues, *in situ* environment, and intrinsic epigenetic mechanisms. Antibodies that recognize rat isotopes for FAPs sorting and isolation via flow cytometry are limited, when compared to those available for mouse and human. VCAM-1 serves as a negative selection marker for FAPs in rat [30]. However, a pro-fibrogenic VCAM-1 positive FAPs population has been reported in mice [40], which, due to technical limitations, we are unable to evaluate in the rat. As more antibodies become available for rat flow cytometry in the future, the development of new flow cytometry panels will enable *in vitro* experimentation on the emerging FAPs subpopulations.

In conclusion, we have identified PGE2 as a novel and critical negative regulator of FAPs fibrogenic and adipogenic differentiation in traumatically denervated skeletal muscle. When considered in the context of the known literature reporting PGE2 as a positive regulator of muscle regeneration and vasculogenesis, our finding here supports a potential future therapeutic role for PGE2 in the management of muscle denervation injuries in human, particularly when long distant nerve transit to muscle reinnervation and the incumbent muscle fibrosis and fat deposition results in permanent physical functional disability.

## CONFLICT OF INTEREST

Christina Doherty, Monika Lodyga, Judy Correa, Caterina Di Ciano-Oliveria, Pamela J Plant, James Bain, and Jane Batt declare that they have no conflicts of interest.

## FUNDING

Funding was provided by Canadian Institutes for Health Research (CIHR) Institute of Musculoskeletal Health and Arthritis (202409MHM-533170-MOV-CEAJ-54987) and 202503MHM-545066-MOV-CEAJ-54987) to Jane Batt and James Bain.

---

**Supplementary Figure 1.** Passaged FAPs Phenotype. FAPs derived from naive gastrocnemius muscle and maintained in cell culture out to 3 passages (P3) demonstrate adipogenic differentiation equivalent to freshly sorted FAPs (P0) as quantified by RT-qPCR for PLIN-1 transcript (A) and immunostaining for PLIN-1 protein expression (B), equivalent fibrogenic differentiation as determined by immunostaining for αSMA protein expression (C), equivalent EP4 receptor transcript expression as determined by RT-qPCR (D) and susceptibility to PGE2 mediated inhibition of adipogenic and fibrogenic differentiation (E), as determined by immunostaining for PLIN-1 (left panel) and αSMA (right panel) protein expression, respectively (Data are mean +/-S.D., ns = not significant [p≥0.05], P = passage number).

**Supplementary Figure 2.** SDS-PAGE and Western blotting for EP4 receptor demonstrate EP4 expression in C2C12 myoblasts, with lesser expression in mature myotubes. Rat FAPs and brain lysate served as positive EP4 receptor controls.

## Notes

### Competing Interest Statement

The authors have declared no competing interest.

