## Supplemental Methods for "Prostaglandin E2 is a Negative Regulator of Fibroadipogenic Progenitor Differentiation in Traumatically Denervated Skeletal Muscle"

**SUPPLEMENTAL MATERIALS**

1. METHODS

*RNA extraction and RT-qPCR*

Flash-frozen gastrocnemius muscle was placed in liquid nitrogen in a RNAse-free mortar and pestle and crushed into a fine powder. The powder was transferred to a 5 mL round-bottom tube on ice, and 1 mL of Trizol (Thermo Fisher 15596026, Burlington, ON, Canada) per 100 mg of tissue was added. Samples were homogenized for 3 × 30 s (Kinematica AG, Lucern, Switzerland) and then processed using a commercially available RNA extraction kit (Invitrogen, 12193555, Burlington, ON, Canada), with on-column DNAse treatment performed according to the manufacturer's specifications (Invitrogen, 12185010, Burlington, ON, Canada).

RNA was isolated from cultured FAPs using a RNeasy Kit (Qiagen, 74104, Germantown, MD, USA), and DNAse treated as per the manufacturer's instructions (ThermoFisher, EN0521, Burlington, ON, Canada).

RNA quantity and quality were determined using an Agilent 2100 Bioanalyzer (Agilent Technologies, Mississauga, ON, Canada). Only RNA with a RIN greater than 7 was used for analyses. 1 μg DNase-treated RNA was reverse transcribed with Superscript III First-Strand Synthesis SuperMix (Invitrogen, 11752250, Burlington, ON, Canada) to generate cDNA. qPCR was performed for genes of interest, with sets of primers designed and validated for specificity and efficiency (Table 1). Gene amplification was performed with Power SYBR green PCR Master Mix (ThermoFisher 4367659, Burlington, ON, Canada) on a QuantStudio7 Flex Real-Time PCR System (ThermoFisher Scientific, Burlington, ON, Canada) with cycling parameters of 95°C for 10 min, followed by 40 cycles of 95°C for 15 s and 60°C for 1 min. Samples were run in triplicate, and a "no-template control" was run for each primer pair. Data were analyzed using the relative comparative delta-delta CT method [Schmittgen, T.D et al. Nat. Protoc. 2008:3,1101–1108]. Hypoxanthine-guanine phosphoribosyltransferase (HPRT), and hydroxymethylbilane synthase (HMBS) served as housekeeping genes.

*Protein Extraction, SDS/PAGE and Western Blotting*

Gastrocnemius muscle was homogenized (Polytron PT 1200E, Kinematica AG, Lucern, Switzerland) in lysis buffer [5 mM Tris/HCl, pH 8.0, 1 mM EDTA, 1 mM EGTA, 1 mM 2-mercaptoethanol, 1% glycerol, PMSF (1 mM), leupeptin (10 μg/mL),aprotinin (10 μg/mL)] three times for 30 s on ice. Homogenates were centrifuged at 1600g for 10 min at 4°C, and supernatant cleared by further centrifugation (10 min, 4 ◦C, 10,009g). Protein lysates were quantified using the Pierce 660 nm Protein Assay Reagent (Thermo Fisher, 22660, Burlington, ON, Canada).

FAPs nuclear and cytoplasmic protein extraction was performed using NE-PER Nuclear and Cytoplasmic Extraction Reagents (Thermo Fisher, 78833, Rockford, IL, USA), as per the manufacturer instructions. To achieve a concentration of at least 1ug/ul of the nuclear fraction (~100ug total nuclear protein), 6-wells of a 6-well plate were pooled per experimental condition prior to commencing the protocol.

Protein lysates were separated on an 8% or 10 % polyacrylamide gel as appropriate, transferred to a nitrocellulose membrane for 1hr at RT, and stained with Ponceau-S (0.1% ponceau, 5% glacial acetic acid) to evaluate protein loading. Blots were blocked in 3% skim milk, blotted with anti-COX-2 (1:1000; ThermoFisher Scientific, PA5-88606, Burlington, ON, Canada)), anti-EP4 (1:1000; Proteintech, 24895-1-AP, Chicago, IL, USA), or anti-PPARγ (1:2500; Proteintech, 16643, Chicago, IL, USA) overnight at 4C and detected with the appropriate HRP-linked secondary antibodies(1:5000) for 1hr at RT. Chemiluminescence was detected using ClarityWestern ECL Substrate (Bio-Rad, 1705060S, Mississauga, ON, Canada) with signal acquired on a Gel Doc EZ Imager (Bio-Rad Laboratories, Mississauga, ON, Canada). Protein expression was quantified on Image Lab 6.0.1 (Bio-Rad, Mississauga, ON, Canada).

*FAPS Immunofluorescence*
Cultured FAPs were fixed with 4% PFA for 15 min at RT, washed with PBS 3 times, and incubated with 100 mM Glycine in PBS for 10min at RT. Cells were washed 3 times with PBS, permeabilized with 0.1% Triton-X in PBS for 20 min, washed twice with PBS, and blocked with 3% Bovine Serum Albumin (BSA) in PBS for 1 hr at RT. Cells were incubated with anti-PLIN-1 (1:400, Abcam, ab3526, Waltham, MA, USA), anti-PPARγ (1:100), anti-Ki67 (1:250, Abcam, AB16667, Waltham, MA, USA), or anti-smooth muscle actin (1:100, Burlington, ON, Canada) overnight at 4◦C and detected with the appropriate secondary antibodies (goat anti-rabbit Alexa Fluor 488 Invitrogen A11008, or Alexa Fluor 555 Invitrogen A21429, or goat anti-mouse Alexa Fluor 555 Invitrogen, A21422, Burlington, ON, Canada) for 1 hr at RT. SMA labelled cells were counterstained with phalloidin iFluor 488 (1:1000, Abcam, AB176753, Waltham, MA, USA). Nuclei were labeled with Hoechst (1:10,000) for 4 min. PBS was added to wells, and plates were stored at 4◦C in the dark until imaged.

*FAPs ORO Staining*
Cultured FAPs were fixed in wells with 10% Neutral Buffered Formalin (NBF, Millipore Sigma HT501128, Oakville, ON, Canada) for 5 min at RT. NBF was removed, and fresh 10% NBF was added for 1h at RT. Cells were washed with 60% isopropanol and allowed to air dry (approximately 2–3 min), followed by incubation with 200 μL of ORO working solution for 10 min at RT. ORO was aspirated, the cells washed 5 times with ddH2O, 1mL of Hematoxylin was added for 3 minutes at RT, and washed an additional 3 times before imaging.

*Script - Fiji Nuclear Count Macro*
```
setOption("ScaleConversions", true);
run("8-bit");
setAutoThreshold("Default dark no-reset");
//run("Threshold...");
//setThreshold(43, 255);
setOption("BlackBackground", true);
run("Convert to Mask");
run("Watershed");
run("Analyze Particles...", "size=100-Infinity display exclude clear include summarize add");
```

2. SAMPLE REFERENCES – UTILIZATION OF TIBIAL NERVE TRANSECTION MODEL

Liu HM, Ferrington DA, Baumann CW, Thomspon LV. Denervation-induced activation of the standard proteasome and immunoproteasome. *PLoS One* 2016 11(11) e0166831.

Liu L, Xiw F, Wei K, Hao XC, Li P, Cao J. Min S. Sepsis induced denervation-like changes in the neuromuscular junction. *J Surg Res* 2016 200 (2) 523-32.

Willand MP, Holmes M, Bain JR, Fahnestock M, De Bruin H. Electrical muscle stimulation after immediate nerve repair reduces muscle atrophy without affecting reinnervation. *Muscle Nerve* 2013 48(2) 219-25.

Yan Y, Sun HH, Hunter DA, Mackinnon SE, Johnson PJ. Efficacy of short term FK506 administration on accelerating nerve regeneration. *Neurorehabil Neural Repair* 2012 26(6) 570-580.

Yohn DC, Miles GB, Rafuse VF, Brownstone RM. Transplanted mouse embryonic stem-cell derived motoneurons from functional motor units and reduce muscle atrophy *J Neurosci* 2008 28(47) 12409-18.

Doherty, C., Lodyga, M., Correa, J., Di Ciano-Oliveira. C. Plant, PJ, Bain, JR, Batt, J. Utilization of the rat tibial nerve transection model to evaluate cellular and molecular mechanisms underpinning denervated-mediated muscle injury. *Int J Mol Sci.* 2024:25(3), pp.1847
