## Supplementary Table 1, FIgures 1 and 2 for "Prostaglandin E2 is a Negative Regulator of Fibroadipogenic Progenitor Differentiation in Traumatically Denervated Skeletal Muscle"

| Gene | Forward Primer | Reverse Primer |
|---|---|---|
| Col1a1 | AAAACGGGAGGGCGAGTGCT | CTCCCTTGGGTCCCTCGACT |
| SMA | CCAGCCAGTCGCCATCAGGA | GCCCGGAGCCATTGTCACAC |
| PLIN-1 | GTGGCTCTCAGCTGCATGT | TTCTGGAAGCACTCACAGGTCC |
| EP1 | TGAGCGCACTGGCCCTCTTG | AGCGCTGGTGATGTGCCGTT |
| EP2 | TGCTCCCTGCCTTTCACAATCTTT | CGCATTAGTCTCAGGACTGGTGGT |
| EP3 | CTTCGAAAGTTCTGCCAGATGATGA | AGGCCGAAAGAAGATACAATCCAAA |
| EP4 | ATTCCGCTCGTGGTGCGAGT | GCCAGAACCACCAATGCGGC |
| 15-PGDH | GGCGCGGCTCAAGGCATAGG | CTGCTCATCCAGGGCCGCTTT |
| PTGS2 | ACGTGTTGACGTCCAGATCA | GGCCCTGGTGTAGTAGGAGA |
| PTGES1 | CAGGCTGCGGAAGAAGGCTTTTG | ATGTCGTTGCGGTGGGCTCT |
| PTGES2 | AAGCGCCTCAAAAGCAGGCAC | ATGGCCGGTCTTTGCCCACG |
| PTGES3 | AGACGGAGCAGATGATGATTCAC | CCCCTCAATATCCAGGCGATG |
| PPARY | GCCTGCGGAAGCCCTTTGGT | AAGCCTGGGCGGTCTCCACT |
| HPRT | GCCGACCGGTTCTGTCAT | TCATAACCTGGTTCATCATCACTAATC |
| HMBS | GGCTCAGATAGCATGCAAGAGA | TGGACCATCTTCTTGCTGAACA |

Supplementary Table 1. RT-qPCR Primers

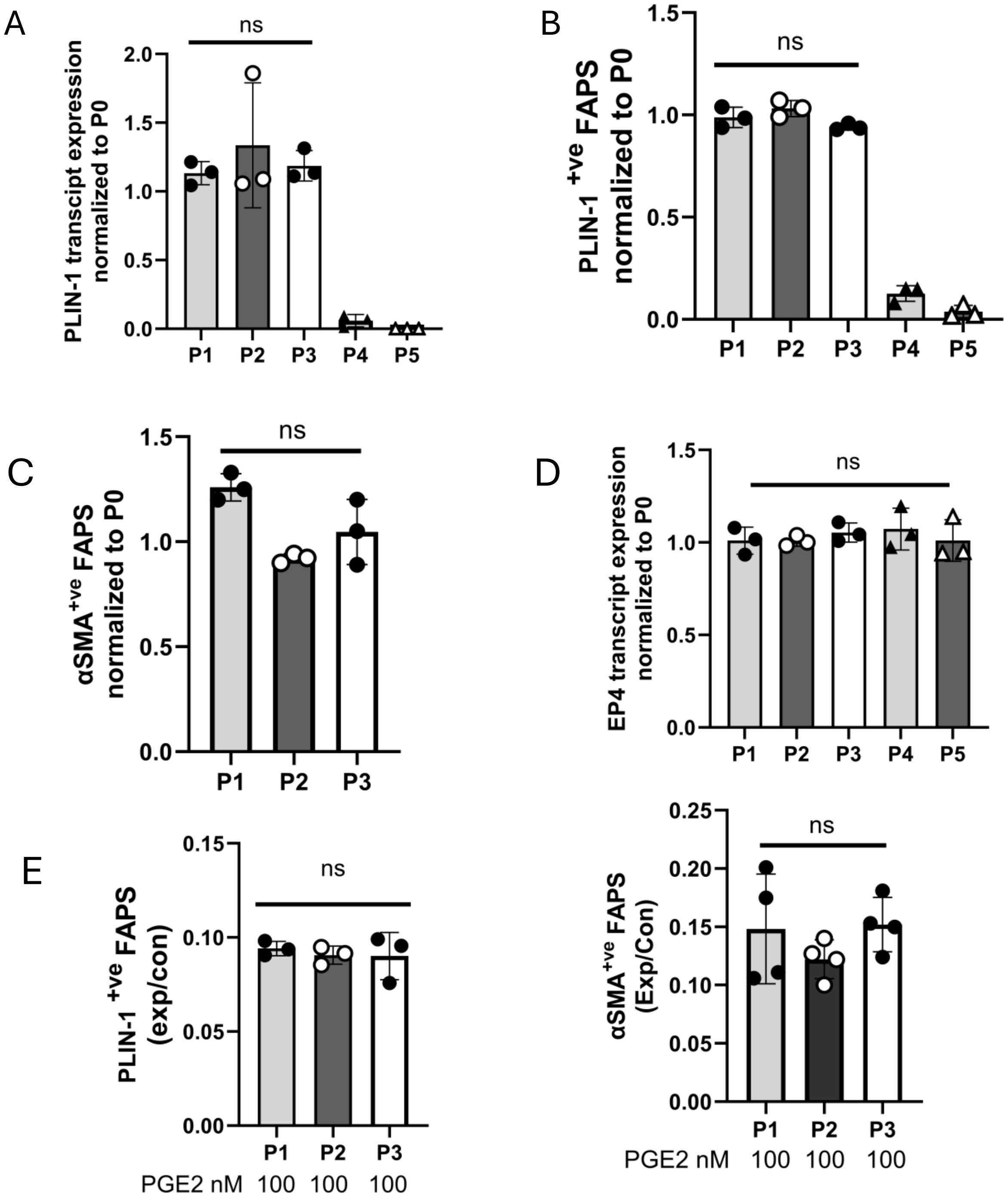


**Supplementary Figure 1**

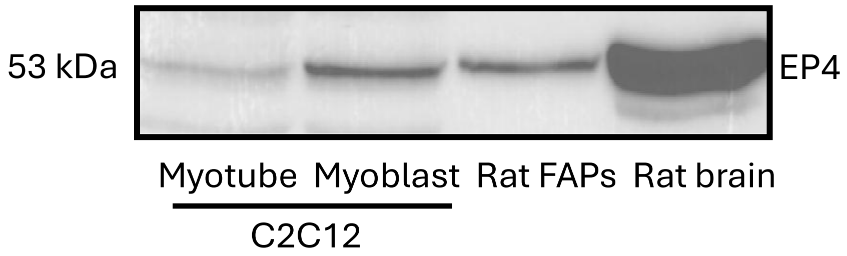


**Supplementary Figure 2.**
